# Explaining Visual Cortex Phenomena using Recursive Cortical Network

**DOI:** 10.1101/380048

**Authors:** Alexander Lavin, J. Swaroop Guntupalli, Miguel Lázaro-Gredilla, Wolfgang Lehrach, Dileep George

**Affiliations:** Vicarious AI, San Francisco, USA

**Keywords:** Visual cortex, Psychophysics, RCN, Bayesian inference

## Abstract

The connectivity and information pathways of visual cortex are well studied, as are observed physiological phenomena, yet a cohesive model for explaining visual cortex processes remains an open problem. For a comprehensive understanding, we need to build models of the visual cortex that are capable of robust real-world performance, while also being able to explain psychophysical and physiological observations. To this end, we demonstrate how the Recursive Cortical Network (George et al., 2017) can be used as a computational model to reproduce and explain subjective contours, neon color spreading, occlusion vs. deletion, and the border-ownership competition phenomena observed in the visual cortex.

## Introduction

For a comprehensive understanding of visual cortex, we need to build models that are capable of robust real-world performance, while also being able to explain psychophysical and physiological observations. One avenue of research considers the tasks of recognition, segmentation, reasoning etc. as queries on a generative model (Lee & Mumford, 2003). Many visual illusions can also be understood as optimal Bayesian inference in a generative model, and they often provide insights into to the mechanisms underlying visual perception. In a recent publication (George et al., 2017), we introduced the Recursive Cortical Network (RCN), a generative model for vision, and demonstrated its real-world performance. Here we show that RCN can reproduce and explain well-known psychophysics experiments and physiological observations: (1) subjective contour effects (Kanizsa, 1976), (2) and neon color spreading, (3) border-ownership response, and (4) occlusion versus deletion effect. All these phenomena are explained as the byproduct of doing inference in the model that was constructed and learned for parsing a visual scene^1^. We argue that these visual phenomena are necessary side effects of the factorizations employed by the model to achieve strong generalization.

## Recursive Cortical Network (RCN)

RCN is a structured probabilistic graphical model (PGM) for vision consisting of a contour hierarchy of features that interacts with an appearance canvas (Fig 1A). The contour hierarchy is learned as alternating layers of feature detectors, pools and lateral connections (Fig 1B). In Figure 1B, each filled circular node is a binary random variable, the open circular nodes are categorical random variables, and the rectangles are factors that encode compatibility. Pooling provides invariance to local deformations, similar to the pooling in convolutional neural nets. The lateral connections, grey square ‘factor nodes’ in Fig 1 B and C, between the pools are learned to enforce contour consistency between the choices in adjacent pools. Figure 1C shows the hierarchical decomposition of a rectangle in terms of simple line segments at the bottom to more complex features in the higher levels, and Figure 1D shows the details of the interactions between contours and surfaces. See George et al., (2017) for details.

**Figure 1:**
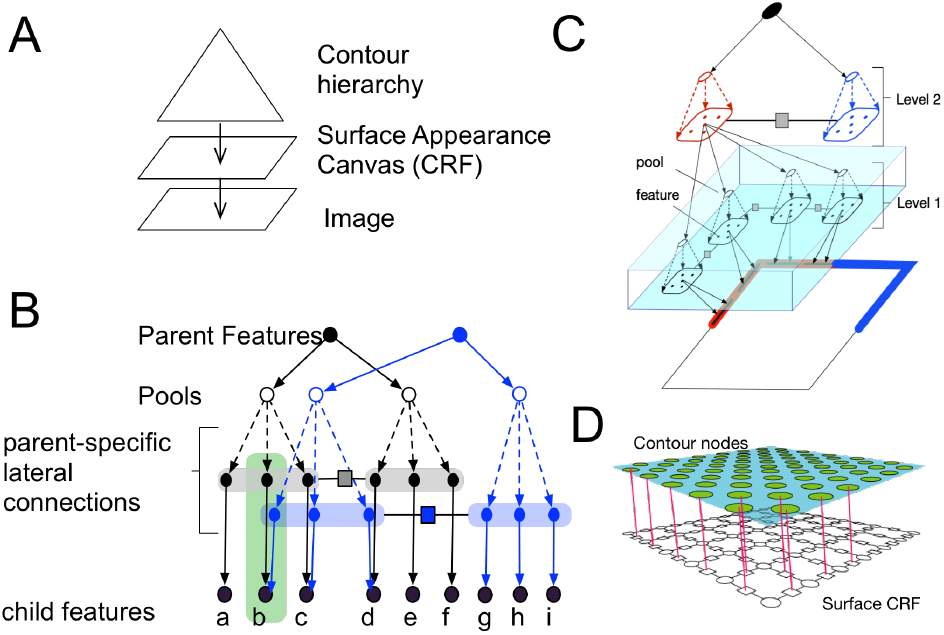
RCN generative model. See text for details.

Parsing a scene is achieved by doing approximate MAP inference (inference to best explanation) using scheduled max-prop belief propagation (Pearl, 1988). The message passing schedule, which was inspired by biology, is as follows. A fast forward pass, which includes short-range lateral propagations, identifies nodes that are highly likely given the evidence. The backward pass focuses on highly active top-level nodes and includes longer range lateral propagations.

The propagations are used to assemble an approximate MAP solution that produces a complete segmentation of the input scene.

## Results

The visual phenomena that we explain share some commonalities. They require the interoperation of feedforward, feedback and lateral connections. Three of them involve the representation of contours and surfaces. All of them can be understood as the result of approximate optimal inference in RCN.

### Subjective Contours

In the subjective contours illusion, people perceive an illusory line that is not supported by local evidence. In Figure 2 top left, people perceive a faint contour of a triangle in the blank space between the circles even though there is no local evidence for a border. Physiological results report evidence for neurons in V1 responding to the illusory contour, albeit with a delay compared to the neurons responding to real contours (Lee & Mumford, 2003). Figure 2 columns 1 & 3 show a diverse of set of images where illusory contours are perceived. In columns 2 & 4, we show how an RCN that is trained to recognize regular shapes ‘hallucinates’ illusory contours in these visual stimuli. What is shown in these images is the ‘inference to best explanation’ (MAP inference) solution at the lowest level of the network, obtained as a result of message passing as described earlier. The yellow portions in these images denote the bottom-up evidence, and the blue stars are the ‘backtraces’ that is part of the global MAP solution found by the network. The backtrace indicates that the network expects to see contours in the blank space to be ON as part of the global solution.

**Figure 2:**
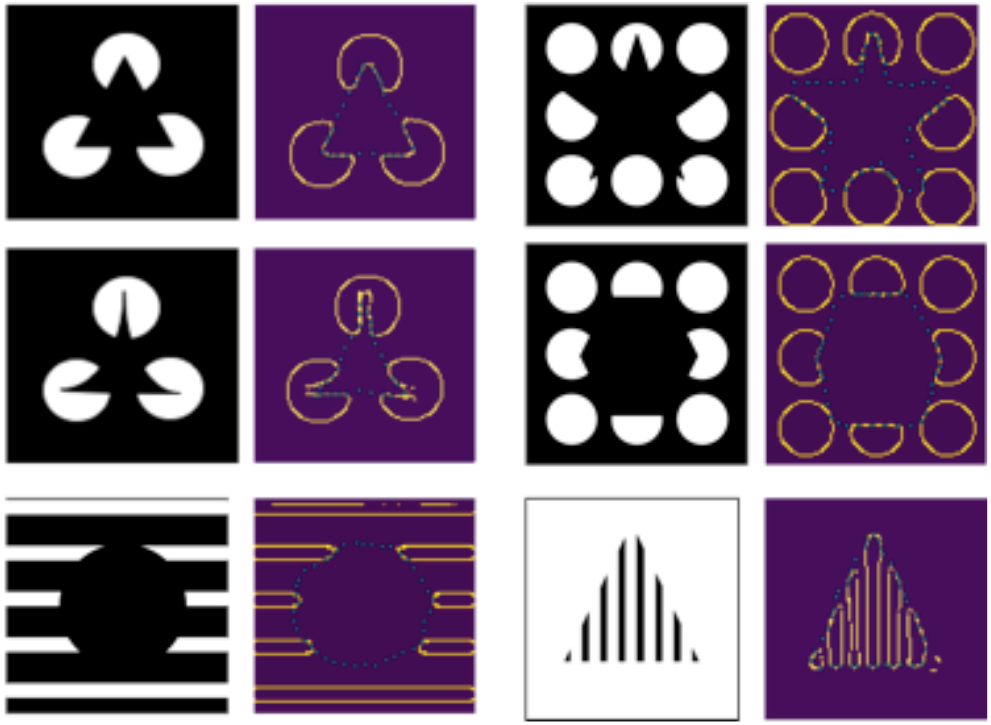
Subjective contours. See text for details.

Why does RCN produce these hallucinations as a result of inference, despite the lack of local evidence? The reason is that the local evidence in the rest of the image is sufficient to support the global percept of the object, according to the model. Since MAP inference finds the configuration that best explains the evidence in the image, it will turn ON all the features that are part of the global percept.

The temporal dynamics of neuronal responses to subjective contours (Lee & Mumford, 2003) can be readily understood from the schedule of message propagation. During the forward pass, the features have only local evidence, and hence the neurons in blank spaces do not respond. Once forward pass identifies a potential global percept, that information flows down in the top-down messages to affect the beliefs in lower level nodes to turn ON some features that were previously OFF.

### Border Ownership Responses

Boundaries of occluding objects are perceived as belonging to them, a property known as *border ownership* (von der Heydt, 2011). Several neurons in V1 and V2 are known to be sensitive to their border ownership. These cells prefer a given figure to be on one side of a border or the other, yet it is not possible to determine from local cues within a cell’s classical receptive field whether a given contour belongs to a surface or not (Tyler 2011). In particular, in the earlier phases of the response to a stimulus, both these copies fire equally, and in the later phase of the response only the neuron with the correct surface selectivity maintain the response (von der Heydt, 2011).

Consistent with findings presented in (von der Heydt, 2011), RCN model has two copies of every contour neuron, one representing each side of the border ownership. However, it is the precise nature of their interaction in the PGM that determines how it arrives at the solution. Figure 3A shows the PGM fragment of RCN corresponding to this interaction. The feature copies with identical contours, but different side-of-surface preferences interact with the contour-node with no preference (‘unselective’) in a noisy-OR ‘V’ structure. The unselective node is directly connected to the rendered image. On the first forward pass through the V-structure, bottom up evidence flows equally to both parents due to lack of prior preference for either of them. Feed forward propagations result in a global percept at the top level that is consistent with only one of the parents. The backward messages then convey these preferences. Since the goal of MAP inference is to ‘explain’ the evidence, one of the parents turning ON ‘explains-away’ the need for the other parent to turn ON. Figure 3B shows the log-likelihoods of the border ownership nodes as a function of the number of message passing iterations. This reproduces and explains the experimentally observed effect.

**Figure 3:**
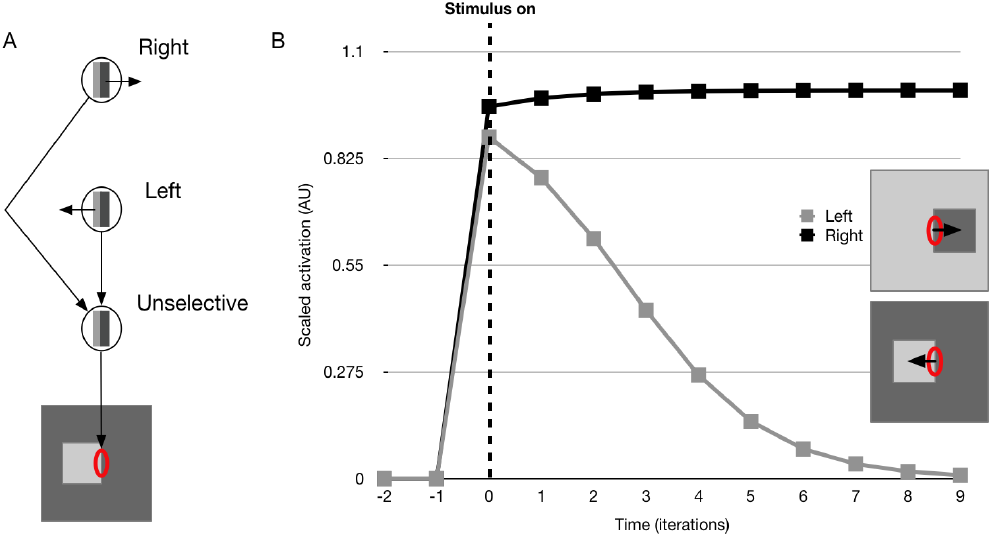
Border ownership experiment with RCN. (A) PGM schematic, (B) Evolution of activation of two contour selective cell copies with identical RFs but opposing border ownership preferences.

### Neon-color Spreading

Certain stimuli, like that in Figure 4 (left), elicit perception of an illusory surface with an illusory color in humans, an effect known as *neon-color spreading* (Bressan et al., 1997). The suggested mechanism behind these effects is the interplay between boundary completion and surface filling-in in visual cortex (Grossberg & Yazdanbakhsh, 2005). Notably, the filling in of the illusory surface respects the boundaries of the illusory contours.

**Figure 4:**
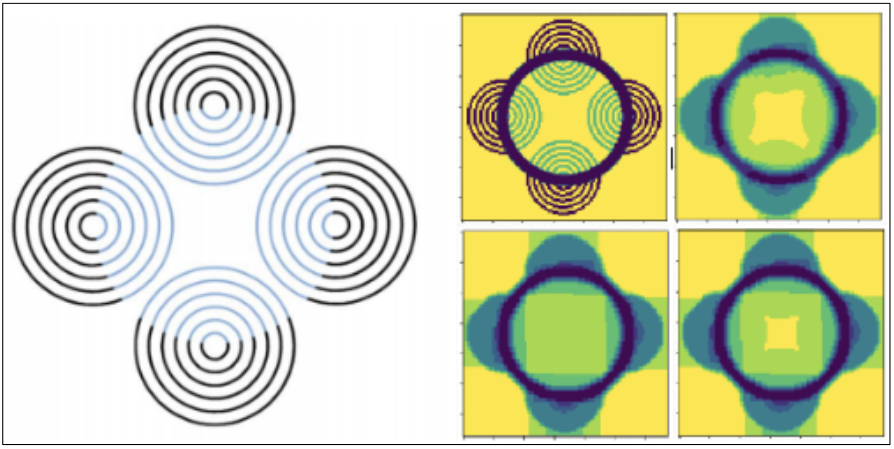
Neon-color spreading experiment with RCN demonstrating the neural filling-in mechanism. Given the input stimulus (Bressan et al., 1997) (left), surface-information is sequentially propagated in the model’s V1 (right, clockwise from top-left).

The neon color spreading effect is a natural byproduct of the dynamics of MAP inference in RCN. To understand this, consider the PGM fragments shown in Figures 1A and 1D. The surface modeled as a conditional random field (CRF) encourages continuity between adjacent surface nodes unless the intervening contour node is turned ON. As described in George et al., (2017), a forward pass through this model produces approximate edge and surface responses. The backward pass, which is based on selecting the most active hypothesis at the top level of the contour hierarchy, will then enforce the corresponding contour discontinuities on the surface CRF. The stimulus shown in Fig 4 (left) has sufficient local edge evidence to support a circle as the top level hypothesis in the contour hierarchy of RCN – this part of the inference is identical to the case of subjective contours described earlier. The top-down partial MAP configuration for contours, the circle, then influences the propagation in the CRF. The discontinuity imposed by the top-down contours will then propagate in the CRF with further message passing to create the fill-in effect.

### Occlusion vs. Deletion

Psychophysics experiments show that humans are much better at detecting objects under occlusion than the same objects with occluded regions deleted (keeping the same visible portion) (Johnson & Olshausen, 2005). In George et al., (2017), we demonstrated that reasoning about occlusions leads to significantly higher recognition rates in RCN.

The reason behind occlusion-vs-deletion is easy to understand in the RCN generative model. Deletion of the parts of an object is absence of evidence for those parts. When those same parts are missing due to occlusion, the model can explain away the absence of evidence as occlusion. Mechanistically, the portions that are deleted will contribute negative evidence to the overall hypothesis if there is no occlusion to explain their absence. Explaining away during occlusion reasoning will convert those negative evidences to ‘uncertain evidence’ (log-likelihood = 0). Figure 5 shows the log likelihood scores obtained for the ‘square’ hypothesis when the missing evidence is treated as occlusion (middle column) vs deletion (right column), in comparison to an intact square (left column).

**Figure 3:**
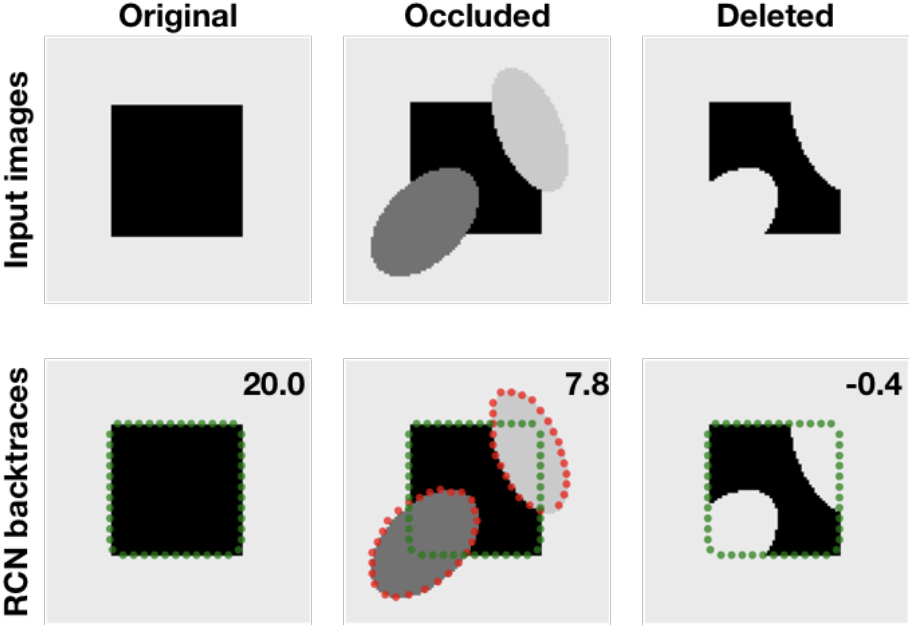
Detection under occlusion versus deletion in RCN. Similar to human psychophysics findings, RCN detection score (top-right corners), reflecting the confidence, is highest when the object is fully visible (left), followed by when it is occluded (middle),

## Discussion

We described how the dynamics of approximate Bayesian inference using loopy belief propagation in RCN could explain several well-known psychophysical and physiological results. In contrast to models that are constructed specifically to explain isolated phenomena, all these observations were explained as the natural byproduct of doing ‘inference to best explanation’ in a model that was learned for parsing a visual scene. Neuro and cognitive science research guided the representational choices and inference algorithms in RCN and those were crucial for it to achieve state of the art performance on several real world benchmarks with very little training data. Our hope is that RCN can be also be used as tool in neuroscience and cognitive science experiments to further understand the computations in visual cortical circuits.

## Footnotes

1 This paper is accepted to CCN 2018. A companion summary paper in the same conference describes the neurobiological mapping of RCN.

## References

Bressan, P., Mingolla, E., Spillmann, L., & Watanabe, T. (1997). Neon colour spreading: a review. Perception, 26, 1353–1366.

George, D., Lehrach, W., Kansky, K., Lázaro-Gredilla, M., Laan, C., Marthi, B., … & Lavin, A. (2017). A generative vision model that trains with high data efficiency and breaks text-based CAPTCHAs. Science, 358(6368)

Gilbert, C. D., & Li, W. (2013) Top-down influences on visual processing. Nat. Rev. Neurosci. 14, 350–363.

Grossberg, S., & Yazdanbakhsh, A. (2005). Laminar cortical dynamics of 3D surface perception: Stratification, transparency, and neon color spreading. Vision Research, 45(13), 1725–1743.

Harrison, W. J., & Rideaux, R. (2017). Voluntary control of illusory contour formation. bioRxiv 219279.

Heydt, R. V., Macuda, T., & Qiu, F. T. (2005). Border-ownership-dependent tilt aftereffect. Journal of the Optical Society of America A, 22(10), 2222.

Huang, X., & Paradiso, M. A. (2008). V1 Response Timing and Surface Filling-In. Journal of Neurophysiology, 100(1), 539–547.

Hubel, D. H., & Wiesel, T. N. (1962). Receptive fields, binocular interaction and functional architecture in the cat’s visual cortex. J. Physiol. 160, 106–154.

Johnson, J. S., & Olshausen, B. A. (2005). The recognition of partially visible natural objects in the presence and absence of their occluders. Vision research, 45(25-26), 3262–3276.

Kanizsa, G. (1976). Subjective Contours. Scientific American, 234(4), 48–52.

Lee, T.S., Mumford, D., (2003). Hierarchical Bayesian Inference in the visual cortex.

Leung, T., & Malik, J. (1998). Contour continuity in region based image segmentation. Computer Vision - ECCV98 Lecture Notes in Computer Science, 544–559.

Marr, D. (1982). Vision: A Computational Investigation into the Human Representation and Processing of Visual Information. San Francisco: W. H. Freeman.

Olshausen, B. A., Anderson, C. H., & Van Essen, D. C. (1993). A neurobiological model of visual attention and invariant pattern recognition based on dynamic routing of information. Journal of Neuroscience, 13(11), 4700–4719.

Pearl, J. (1988). Probabilistic Reasoning in Intelligent Systems: Networks of Plausible Inference. Morgan Kaufmann.

Zhou, H., Friedman, H. S., & Heydt, R. V. (2000). Coding of Border Ownership in Monkey Visual Cortex. J. Neurosci, 20(17), 6594–6611.

